# *In silico* discovery of repetitive elements as key sequence determinants of 3D genome folding

**DOI:** 10.1101/2022.08.11.503410

**Authors:** Laura M. Gunsalus, Michael J. Keiser, Katherine S. Pollard

## Abstract

Natural and experimental genetic variants can modify DNA loops and insulating boundaries to tune transcription, but it is unknown how sequence perturbations affect chromatin organization genome-wide. We developed an *in silico* deep-learning strategy to quantify the effect of any insertion, deletion, inversion, or substitution on chromatin contacts and systematically scored millions of synthetic variants. While most genetic manipulations have little impact, regions with CTCF motifs and active transcription are highly sensitive, as expected. However, our analysis also points to noncoding RNA genes and several families of repetitive elements as CTCF motif-free DNA sequences with particularly large effects on nearby chromatin interactions, sometimes exceeding the effects of CTCF sites and explaining interactions that lack CTCF. We anticipate that our available disruption tracks may be of broad interest and utility as a measure of 3D genome sensitivity and our computational strategies may serve as a template for biological inquiry with deep learning.

## Introduction

The human genome gives rise to its own organization in the nucleus, where the folding of chromatin into intricate and hierarchical structures can be reflective and instructive of cell state [1]. Sequence itself contains the information to create some chromatin features. Binding of CTCF proteins to DNA motifs blocks the extrusion of DNA by motor proteins to create topologically associating domains (TADs) spanning hundreds of megabases [2–5]. These dynamic structures permit interaction of elements within their boundaries and limit interaction with elements outside to tune gene expression [6,7]. However, recent reports reveal CTCF may not be the only factor involved, as some contacts remain after CTCF depletion, and interactions across megabases are not affected [8,9]. How exactly sequence informs structure ranging from the highest levels of genome organization—chromosome territories and compartments—to the level of individual enhancer-promoter interactions, still remains unclear.

Current approaches relating genome sequence to folding either leverage natural genetic variation or experimentally manipulate particular loci to test specific hypotheses. Applying chromatin capture to genetically diverse individuals revealed single nucleotide variants associated with loss or gain of chromatin contact [10]. Large structural variants are also rare at domain boundaries in healthy humans but not in patients with autism or developmental delay [11]. To understand the mechanisms underlying these associations, experimental studies have engineered chromatin contact in cells and mice with synthetic tethering [12] and CRISPR systems [13–15] and measured their effects on genome folding and expression of genes such as *Hbb* and *Vcan*. Findings in these individual loci may not apply genome-wide and could overlook mechanisms without known precedent. Here, we propose combining the genome-wide power of population genetics with the precision seen in experimental studies. We develop a strategy which leverages deep learning to comprehensively screen the human genome for key regulators of 3D genome folding.

Whereas previous machine learning approaches required domain experts to select the most relevant features, deep learning allows patterns to be learned directly from the data without expert input. Deep learning models perform well in predicting enhancer activity [16,17], transcription factor binding [18], gene expression [19], and genome folding [20,21] from sequence, with newer models increasing scale and incorporating epigenetic assays to provide cell type-specific context [22–24]. We can probe these models as computational oracles to predict the behavior of DNA sequence at scales intractable experimentally [25]. Models have been applied to predict the impact of structural variants on human genome folding [20,26], confirm the importance of CTCF through computational mutagenesis [20], and resurrect the folding of Neanderthal genomes [27]. These early reports show that many highly disruptive perturbations lack CTCF or annotated regulatory elements, hinting that there may be sequences that encode information needed for genome folding left to uncover.

Here, we leverage Akita [20], a convolutional neural network trained to predict genome folding from sequence, to develop a computational method that performs unbiased and targeted *in silico* mutagenesis experiments at scale. Applying this approach to a human foreskin fibroblast cell line (HFFc6) with high-resolution micro-C data for model training, we discover wide variability in how robust genome folding is to sequence perturbations across loci. Investigation of sensitive loci reveals both known motifs, like CTCF, and understudied modulators of 3D genome folding, including transposon and RNA gene clusters. Our genome-wide screen reveals a diverse vocabulary of DNA elements that collaborate with CTCF to orchestrate TAD-scale chromatin organization.

## Results

### Genome-wide deletion screen reveals high variability in 3D genome folding

To measure sequence importance to chromatin organization, we developed a deep-learning scoring strategy to computationally introduce modifications into the human reference genome and predict their impact on genome folding with Akita [20]. Given a ∼1-megabase (Mb) DNA sequence, this model accurately produces a chromatin contact map at ∼2-kilobase (kb) resolution, where TADs and DNA loops are visible. We quantify the impact of a centered sequence variant, which we call *disruption*, as the log mean squared difference between the predicted contact frequency map for the 1-Mb sequence with a sequence alteration compared to that of the reference sequence. If a variant dramatically rearranges how the genome is predicted to fold, we infer that the original sequence could regulate chromatin contacts. In this study, we perform a variety of genome-wide sensitivity screens across millions of genetic perturbations, including targeted and unbiased deletions, insertions, and substitutions ranging from 1 base pair (bp) to 500,000 bp (**Fig. 1a**). In contrast to *in vivo* genetic perturbations, our approach enables precise and flexible genome editing at scale.

**Figure 1:**
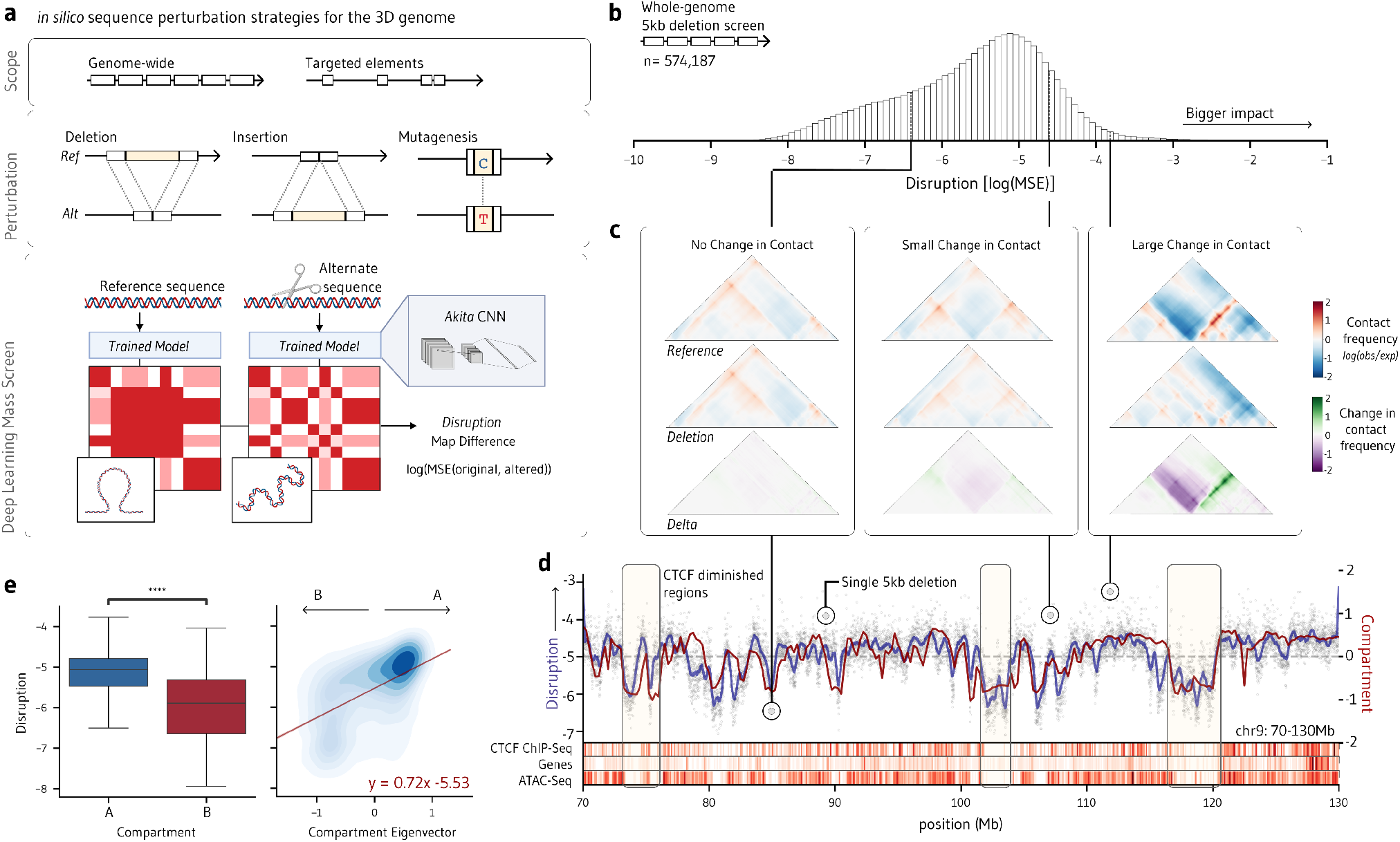
*In silico* deletion screen indicates the impact of sequence perturbation on 3D genome folding is highly variable. **a**. We quantify how important DNA sequence is to genome folding by introducing whole-genome and targeted deletions, insertions, and point mutations and comparing the predicted Hi-C contact maps to maps predicted from the reference sequence. We score *disruption* as the log mean squared difference of the perturbed map relative to the reference map (MSE). Variants with high disruption scores are inferred to contribute to 3D genome folding. **b**. A genome-wide tiled 5-kb deletion screen produces a distribution of sequence importance with log(MSE) between -10 and -1 for the HFFc6 cell type. **c**. Genome-wide screens capture a range of disruption scores; some sequences do not change predicted genome folding (left panel), some produce small focal changes (middle panel), and others dramatically rearrange boundaries (right panel). **d**. The rolling average of disruption and compartment score across a 60-Mb region of chromosome 4. Peaks correspond to regions sensitive to perturbation, while valleys indicate regions robust to perturbation. Yellow shading highlights genomic regions with relatively few CTCF motifs. These regions have low disruption scores, suggesting that their perturbation has little effect on genome folding. **e**. Sensitivity to disruption correlates strongly with compartment score, as measured by the first eigenvector of HFFc6 micro-C.

We first leveraged this strategy to assess all 5-kb deletions tiled across the genome for their impact on folding in HFFc6 cells (n=574,187). Deletions are highly variable, and around half produce changes to chromatin contact maps that are noticeable by eye (**Fig. 1b**). Some sequence deletions completely rearrange the boundary structure of contact maps, some result in small focal changes (e.g., gain or loss of a loop anchor), and some produce no change at all, suggesting the chromatin structure is robust to sequence manipulation (**Fig. 1c)**. As expected, regions of the genome with many CTCF motifs are particularly sensitive while regions with no motifs are perturbation-resilient (**Fig. 1d**). In sum, 62.1% of the most sensitive sequences (top decile of scores) fall within 5 kb of CTCF-bound distal enhancers, compared to only 7.3% of the most robust sequences (bottom decile of scores), establishing that our approach identifies known genome folding mechanisms (**Fig. 2a**).

**Figure 2:**
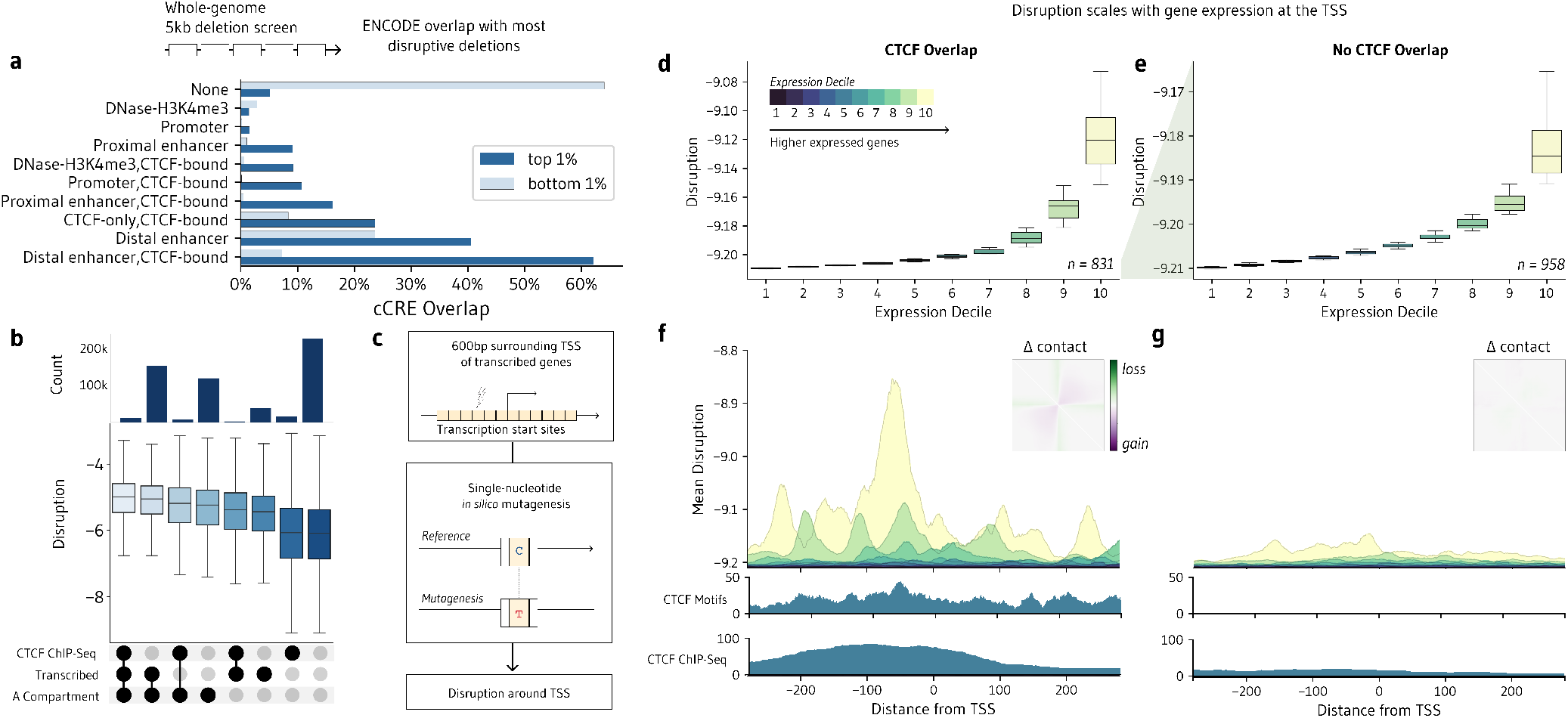
Transcription and CTCF are key modulators of 3D genome folding. **a**. Overlap between top 1% (most disruptive; dark blue) or bottom 1% (least disruptive; light blue) 5-kb sequence deletions and ENCODE candidate cis-regulatory elements, quantified as the proportion of deletions with overlap. Each deletion may overlap with more than one regulatory element. **b**. Average disruption score across genomic regions overlapping with CTCF ChIP-seq peaks, A compartments, and/or actively transcribed sequences. **c**. Single base-pair mutagenesis screen of a 600-bp segment surrounding the transcription start site (TSS) of the most highly transcribed genes in HFFc6 (n=1,789). **d-e**. Mean disruption score of transcribed genes, stratified by expression level decile (colors), and separated into those whose TSS region overlaps (**d**) versus does not overlap (**e**) with CTCF sites. **f-g**. Average disruption score of each base at TSS regions with (**f**) and without (**g**) a CTCF motif overlap, stratified by expression decile (colors), along with average CTCF motif density and CTCF ChIP-seq. Metaplots (upper right) show the average change in contact for the 100 TSSs with the most significant disruption scores.

### Perturbing epigenetically active regions disrupts genome folding

Disruption scores are also correlated with chromatin compartment, as measured by the first eigenvector of the experimental HFFc6 micro-C contact matrix (Pearson’s *r* = 0.522, *P* < 1 × 10^−300^, *n* = 11,413; **Fig. 1e**) [28]. The mean disruption score within gene-rich and open A compartments is 14.6% higher than in compact, inactive B compartments. High gene density and GC content are associated with peaks in disruption as well (**Fig. 1d, Fig. S1a-c**). Using HFFc6 total RNA-Seq [29], we quantified transcription in each 5-kb window and observed a strong correlation with disruption scores (Pearson’s *r* = 0.366, *P* < 1 × 10^−300^, *n* = 11,413). Other genomic features associated with active chromatin are also more frequent in the most sensitive sequences, including distal and proximal enhancers and promoters (**Fig. 2a**). In sum, it is difficult to perturb inactive chromatin and easy to perturb active chromatin.

The correlation between many of these features reflects an inherent challenge in disentangling which are causal and which are reflective of genome folding (**Fig. S1c**). Indeed, regions that are in A compartments, contain CTCF binding sites, and are actively transcribed are also the most sensitive (**Fig. 2b**). The effect of CTCF holds within both A and B compartments (**Fig. S2**), indicating that it is directly associated with sensitive 5-kb bins and is not just a proxy for A compartments. However, both transcription and compartment are more impactful individually than the presence of CTCF motifs, suggesting additional rules govern which CTCF sites are in use and which are redundant or decommissioned. Overall, our findings suggest that independent mechanisms at epigenetically active regions may collaborate to coordinate genome folding.

### Transcriptionally active regions modulate folding alongside CTCF

Elevated disruption scores in gene-dense A compartments motivated us to carefully investigate transcription start sites (TSS). CTCF binding is essential for activity of some promoters [8], and emerging work reveals RNA polymerase II and transcription may separately influence 3D genome folding [30,31]. To test this hypothesis, we evaluated all single-nucleotide mutations in the 300 base pairs (bp) on either side of the TSS of the 1,789 highest expressed protein coding genes in HFFc6 [29] and compared disruption scores to expression level in regions where CTCF motifs are present or absent (**Fig. 2c**). Regardless of CTCF, disruption scales with gene expression (**Fig. 2d,e**). In regions flanking a CTCF motif, we observe a strong peak in disruption directly upstream of the TSS (**Fig. 2f, Fig. S3**). The periodic pattern is more detailed than underlying CTCF motifs and more precise than a sum of CTCF ChIP-Seq peaks around the TSS. Metaplots of the average change in contact reveal that mutations weaken boundaries at the TSS. Our analysis points to a presence of CTCF at the promoters of highly expressed genes, where some CTCF motifs are selectively bound and some are not. We note that even when no CTCF is present, disruption is still slightly elevated upstream of the TSS of highly transcribed genes (**Fig. 2g, Fig. S3**). Active transcription may provide an alternate means of stabilizing DNA-DNA interactions in TSS devoid of CTCF sites through uncharacterized mechanisms, like transcriptional machinery, nascent RNA, or recruited regulatory RNA.

### Transposon clusters modulate genome folding independently of CTCF

At the chromosome scale, we observed clusters of Alu elements and some other repetitive elements alongside peaks in disruption scores, motivating us to explore their role in 3D genome folding (**Fig. 3a**). DNA and RNA transposons replicate and insert themselves into DNA, and constitute over 50% of the human genome [32,33]. They are rich in transcription factor binding sites [34–36], suggesting that some may have been evolutionarily repurposed as regulatory elements. Growing evidence indicates they provide a source of CTCF motifs across the genome and serve as both loop anchors and insulators [37–39]. To measure the impact of different families of repetitive elements on 3D genome sensitivity, we compared disruption of 5-kb windows containing repetitive elements to those with none. Several families exhibit greater sensitivity to perturbation than CTCF containing regions (e.g., Alu, SVA, scRNA, srpRNA; **Fig. 3b**). Disruption scores of repetitive elements are not correlated with mappability, indicating that poor micro-C read mapping in model training data does not bias this result (**Fig. S4, Supplemental Note**). As with CTCF [40], regions with higher numbers of Alu elements are more disruptive upon deletion: the disruption score of 5-kb windows with 5 or more Alu elements is 9.88% higher than that of windows with no elements (*P* < 1.54 × 10^−291^; **Fig. 3c**). This clustering effect holds across many repetitive elements, including MIR and L2 LINE elements, as well as across most small, non-coding RNA genes (**Fig. 3c**). Many families, like L1 LINE elements, show no correlation at all, and trends are consistent across both A and B compartments, hinting that clustering is family specific (**Fig. 3c, Fig. S5**).

**Figure 3:**
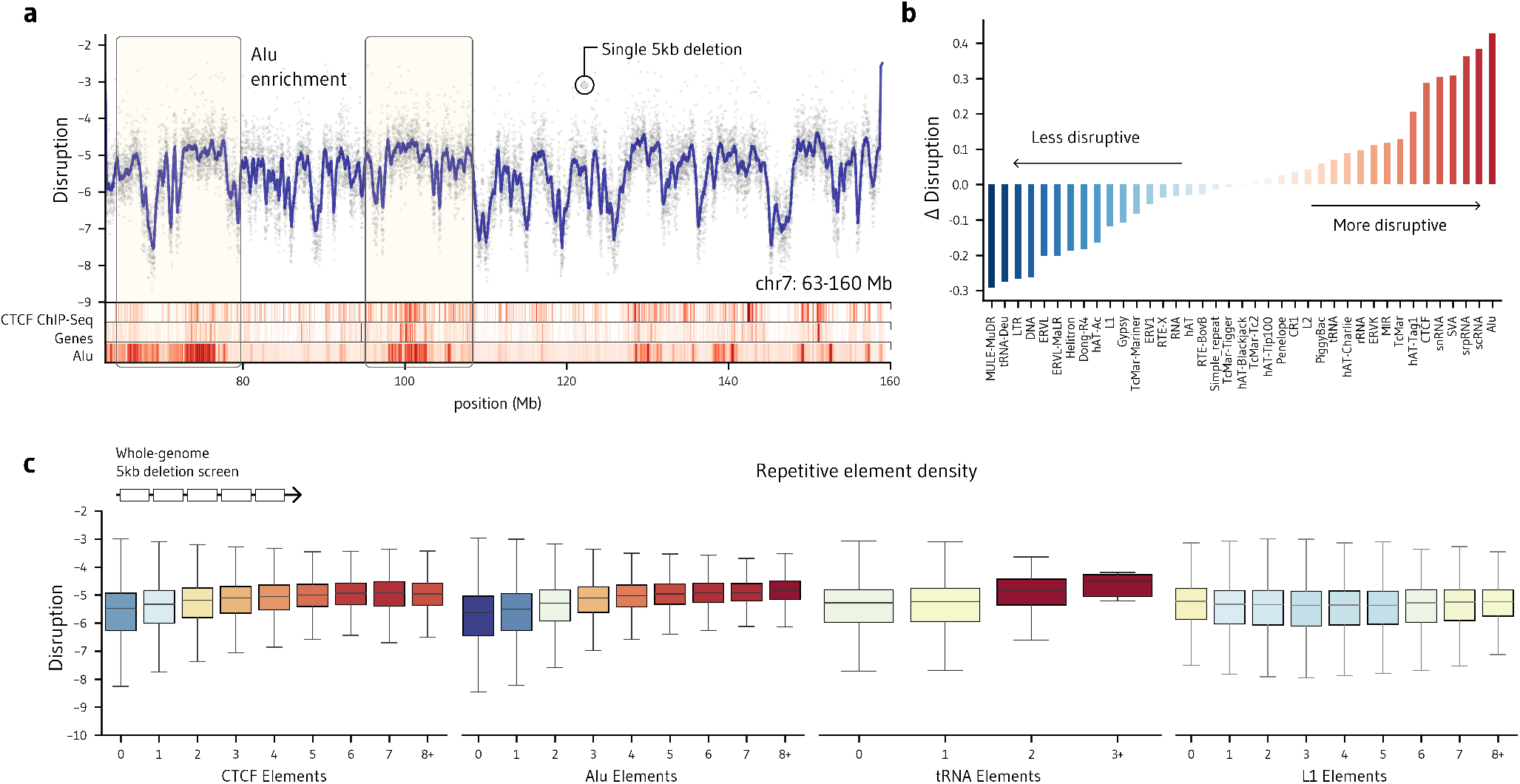
Regions with repetitive elements are sensitive to sequence perturbation. **a**. Mean disruption scores of tiled 5-kb deletions across a 100-Mb region of chromosome 7. Tracks below the plot illustrate the density of CTCF motifs, genes, and Alu elements. **b**. Mean difference in disruption scores between windows containing at least one repetitive element and windows containing none, stratified by family. **c**. Disruption scores of 5-kb deletions stratified by the number of Alu elements, tRNAs, L1 LINE elements, and CTCF motifs they contain.

To investigate the contribution of repetitive elements independently of flanking sequence, we next individually deleted over 1 million elements in the RepeatMasker interspersed repeat database (**Fig. 4a**). Overall, many elements create large-scale boundary shifts, with some causing increases and others decreases in contact frequency (**Fig. 4b)**. Deletions of almost all families are more disruptive than random deletions, and deletions of families such as Alu, small RNAs, SVA, and hAT-Charlie are on par with or exceed deletions of CTCF sites across the genome (**Fig. 4c**). Disruption is moderately correlated with size, but many highly disruptive element families are relatively small and cause unexpectedly large disruptions given their length (**Fig. S1d-e, Fig. 4c**). For example, deletion of tRNAs, scRNAs, srpRNAs, and snRNAs–all under 130 bp on average–drastically alter genome folding on average.

**Figure 4:**
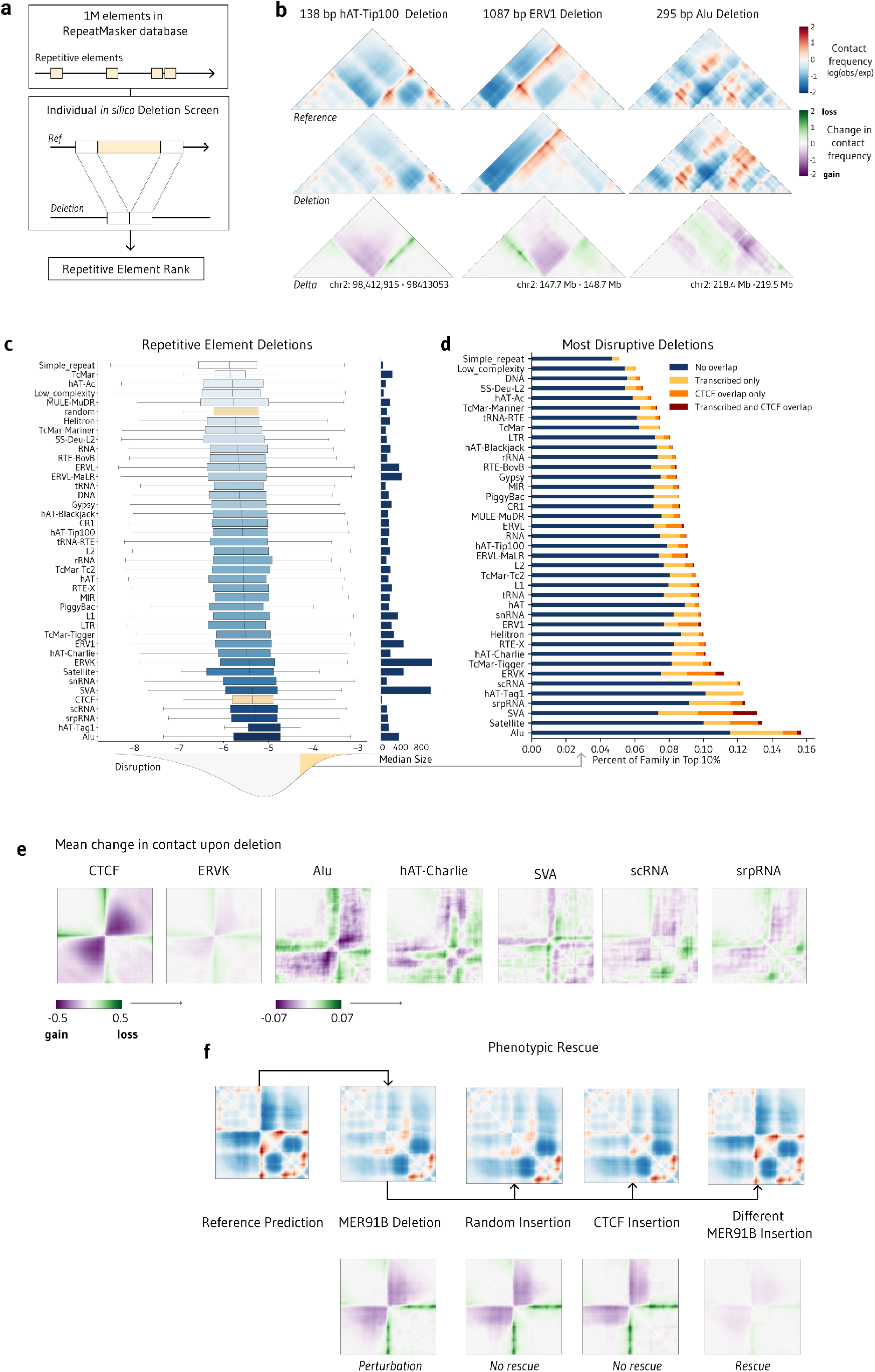
Repetitive element deletions impact genome folding. **a**. Strategy to individually delete over 1 million elements from the RepeatMasker database. **b**. Representative examples from chromosome 2 showing how the deletion of a hAT-Tip100 element, an ERV1 element and an Alu element *in silico* significantly alter contact maps. Single elements are predicted to disrupt genome folding. **c**. Distribution of disruption scores across each repetitive element family (n = 1,164,108). The distribution of disruptions from 100,000 CTCF deletions (positive control) and 100,000 100-bp random deletions (negative control) are shown in yellow. The median size in base pairs of deleted elements for each family is shown on the right. **d**. The top 10% most disruptive elements across the screen by repetitive element family. Most elements do not overlap a CTCF motif or a region actively transcribed in the HFFc6 cell line. **e**. Average changes in contact maps for the top 100 elements per family. **f**. Phenotypic rescue. We showcase a 138-bp MER91B hAT-Tip100 element whose deletion produces a loss of a boundary. Inserting a random size-matched sequence and a CTCF motif does not change the disturbed contact map, but introducing an MER91B element from the same family restores the original genome folding.

We next explored possible mechanisms of repetitive elements in genome folding. Causality is challenging to untangle since each repetitive element can contain features with known associations to chromatin organization. First, the lengths of repeat clusters are roughly similar to clusters of CTCF motifs at TAD boundaries (**Fig. 4c**). Second, several repeat families are known to harbor CTCF motifs [41]. Third, some repeats have a strong GC bias (e.g., Alu GC% > 50%), potentially allowing them to establish compartments [42,43]. Finally, repetitive elements collectively account for a large amount of total nuclear transcription [32]. To dissect the contributions of CTCF and active transcription versus other features of repetitive elements, we quantified overlap of these two annotations with repetitive elements with the highest disruption scores. Only 5.86% of the 10% most disruptive elements contain a CTCF motif while 13.55% are actively transcribed (**Fig. 4d**), so a majority overlap neither. Disruptive repetitive element deletions are enriched at distal enhancers that are not CTCF bound (**Fig. S5d**). These findings hint that repetitive elements may aid in chromatin loop extrusion independently and in collaboration with CTCF and transcription.

To understand the folding phenotypes of element deletions, we next averaged the changes in contact frequency for the top-scoring elements of each family (**Fig. 4e**). ERVK elements behaved like CTCF sites: their deletion led to a strong and centered loss of a chromatin boundary. Other repeat families created an off-diagonal gain in contact, as seen with Alu and hAT-Charlie, dispersed focal disruption, as with non-coding RNAs, and stripes, as with SVA elements. To demonstrate that the model is internally consistent, we performed a *phenotypic rescue*, where we deleted an individual hAT-Tip100 element to produce a large change in contact and attempted to restore the original folding pattern with a different sequence (**Fig. 4f**). While introducing random DNA or a CTCF motif did not recreate the original contact, inserting a related MER91B hAT-Tip100 element did. We conclude that repetitive element families are associated with distinct chromatin contact map features, and elements within a family are functionally interchangeable.

### Insertion of repetitive elements leads to distinct folding phenotypes

Our deletion experiments do not distinguish between repetitive elements that collaborate with CTCF to weaken or strengthen nearby TAD boundaries and those that separately create chromatin contact. To isolate the effects of repetitive elements, we next designed *in silico* insertion experiments. We first engineered a “blank canvas” with no predicted structure by depleting a randomly generated 1 Mb DNA sequence of all CTCF-like motifs (**Fig. 5a, Fig. S6**). We then developed a pipeline to insert one or more copies of any query sequence into this 1 Mb and quantify newly arising chromatin contacts. We easily recreated a division closely resembling a TAD boundary by inserting multiple copies of the canonical CTCF motif (**Fig. 5b**), validating this approach in creating chromatin contact phenotypes.

**Figure 5:**
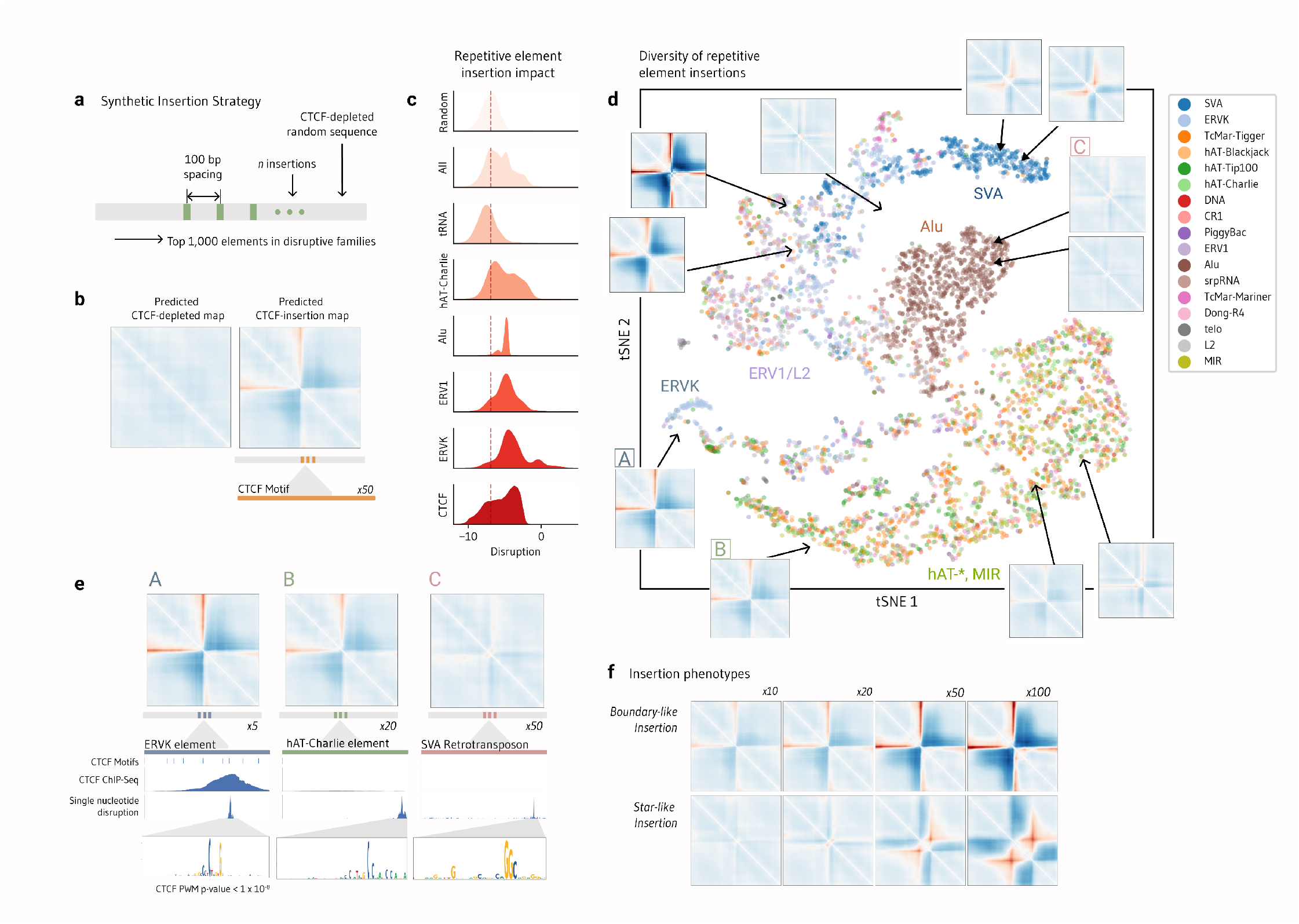
*In silico* insertion screen reveals repetitive elements can induce different boundary types. **a**. Insertion screen strategy. For each of the 1,000 most disruptive elements, up to 100 individual copies (green) are inserted 100 bp apart centered in a 1-Mb random DNA sequence depleted of CTCF sites. **b**. The map predicted from the CTCF-depleted random sequence (left panel) provides a blank canvas against which we can measure the impact of insertions. A CTCF site insertion into the middle of the sequence produces boundaries in the predicted maps (right panel). Disruption is measured as the mean squared difference between the blank map and the predicted post-insertion map. **c**. Distribution of disruption scores across repetitive element insertions (n = 14,514). The score distributions of 10,000 100-bp random insertions (negative control) and of 10,000 CTCF motif insertions (positive control) are shown. **d**. t-SNE visualization of all predicted maps from repetitive element insertions with an disruption score above -5.5. Predicted maps are colored by element family. **e**. We highlight three repetitive elements which are highly disruptive both when deleted and inserted. We overlay overlapping annotated CTCF motifs and CTCF sites confirmed by ChIP-Seq in HFFc6 cells. We also show the disruption score of each nucleotide across the element following single base pair *in silico* mutagenesis, highlighting the motif within the repetitive element responsible for the element’s high disruption score. **f**. We observe two primary classes of insertions: CTCF-like boundary insertions are common across ERVK and ERV1 elements and star-like insertions are common across SVA and Alu elements.

After introducing the 1,000 most disruptive repetitive elements in our deletion screen into a blank canvas, we find a majority also changed contact with insertion, including 80.3% of Alu elements and 86.0% of ERVK elements (**Fig. 5c**). Additional copies strengthened impact, and fewer copies were needed to induce a chromatin boundary compared to the CTCF motif (**Fig. S7)**. Clustering the insertion maps revealed hAT/MIR insertions produced distinct folding patterns from ERV/SVA element insertions (**Fig. 5d**). Alu elements consistently produced focal changes at the site of insertion that appear unlike CTCF-like boundaries. Curiously, repetitive elements seem to produce two distinct modifications to 3D structure upon insertion. Some elements create CTCF-like domain boundaries which increase in strength as more elements are inserted (**Fig. 5f**). Other elements, like the Alu family, create star-like partitions which increase in size with more element insertions. Insertions of tRNA genes did not create new boundaries, suggesting that their effect on 3D genome folding may be context dependent.

Some repetitive elements harbor CTCF motifs and overlap with CTCF ChIP-Seq peaks, strongly suggesting that the model predicted their importance because they contain CTCF binding sites. To test this hypothesis, we performed saturation mutagenesis across a number of high scoring repetitive elements (Fig. 5e). Screening an ERVK element, for example, revealed that the single nucleotides predicted to have the highest importance for contacts lie directly at the center of a CTCF binding site (Fig. 5e). Overall, the closer a sequence is to matching the canonical CTCF motif, the larger the predicted impact of its insertion (Fig. S8). Still, most of the elements that produced contact changes had no CTCF overlap, and the 5 to 50-bp motifs within these elements with the greatest impact did not resemble CTCF motifs (Fig. 5e, Fig. S9). Therefore, insertions support the hypothesis that repetitive elements contain sequence determinants of 3D genome folding beyond CTCF motifs.

### Necessary vs Sufficient: A 60 bp segment of Charlie7 is sufficient to induce a CTCF-like boundary

Mutating individual nucleotides can be enough to disturb protein binding and profoundly impair 3D folding. By contrast, creating a boundary, loop, or domain from scratch is more challenging, and it is fundamentally unclear what minimum sequence is sufficient. We next extended our screening approach to explore which subsequences can produce the *de novo* contact of a full element.

First, we examined CTCF motifs. Fudenberg et al. mutated all motifs in the JASPAR transcription factor database and determined that CTCF and CTCFL are most sensitive to sequence perturbation [20]. To complement this work, we inserted all motifs into a blank map. We find that CTCF and CTCFL are the transcription factor motifs best able to induce genome folding independently of any surrounding genomic context, followed by HAND2, Ptf1A, and YY2 (**Fig. 6a**). Sampling and inserting motifs from the CTCF position weight matrix, we found that the consensus sequence creates a stronger boundary than 99.50% of CTCF variants (**Fig. 6b**). A small minority of CTCF super-motifs with a T at positions 8 and 12 outperformed the canonical motif, hinting that the most commonly bound CTCF motifs may not be the most strongly insulating ones.

**Figure 6:**
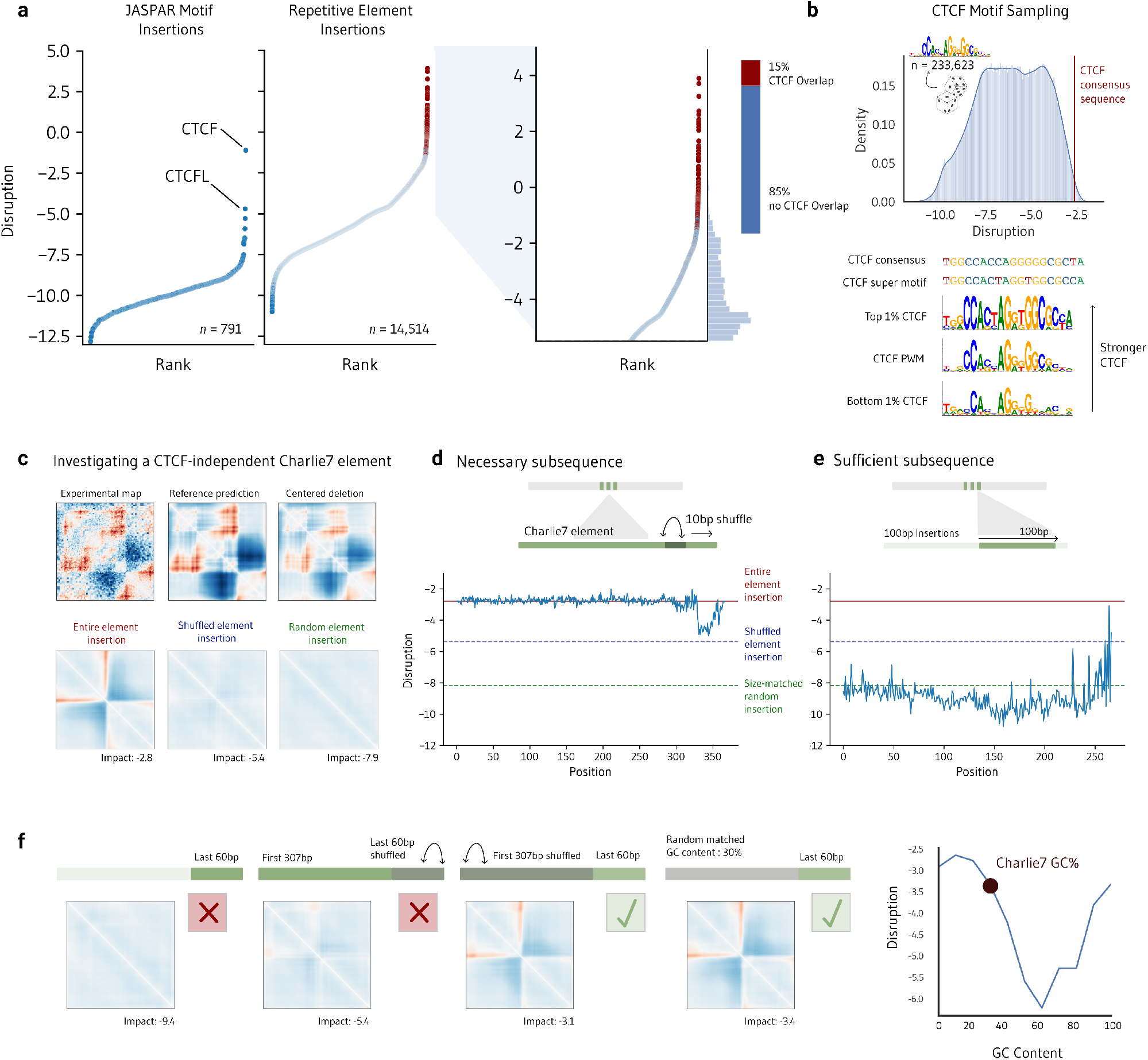
*In silico* investigation of sequence features necessary and sufficient for repetitive element Charlie7 to create a boundary. **a**. We insert every JASPAR motif into a CTCF-depleted random sequence, as well as 14,514 repetitive elements, and rank them according to their disruption score. 85% of the most impactful insertions (score > -5.5) do not overlap a CTCF motif. **b**. We generate CTCF motif variants with frequencies sampled from the CTCF motif position weight matrix (PWM) and insert them into the random reference sequence (n = 326,177), finding that 0.50% of motifs produce stronger predicted boundaries when inserted than the CTCF consensus sequence. These ‘super motifs’ share Ts at positions 8 and 12. **c**. We investigate a 367-bp disruptive Charlie7 hAT-Charlie element which does not overlap a CTCF motif or ChIP-Seq peak. Shown in the top row are the experimental micro-C contact map around the locus of the Charlie7 insertion, the map of the locus predicted by Akita, and the predicted map following the deletion of the entire element. Shown in the bottom row are the predicted maps after insertion into the reference, CTCF-depleted sequence of the Charlie7 element (left), a version of the element with a shuffled sequence (middle) and a random sequence of equal length (right). **d**. We shuffle each 10-bp subsequence along the element to determine which one is necessary to produce the boundary seen from introducing the whole element. **e**. We introduce 100-bp segments scanning the entire element into the reference sequence and find that none is sufficient to produce a strong boundary. **f**. A DNA sequence matching the GC content of Charlie7’s first 307 bp combined with the last 60 bp is sufficient to recreate a boundary. Right panel: The first 307 bp of Charlie7 was replaced with randomly generated sequence across a range of GC content.

Next we dissected Charlie 7, a 367-bp AT-rich (29% GC) hAT-Charlie element on chromosome 11. Deleting Charlie 7 eliminates chromatin interactions (**Fig. 5c; Fig. 6c**). Inserting twenty tandem copies of Charlie 7 creates a CTCF-like boundary, despite no subsequence resembling a CTCF motif. This boundary could not be reproduced by inserting a shuffled Charlie7 sequence or a random sequence of the same length. We therefore shuffled individual 10-bp segments of Charlie 7 to destroy local sequence grammar before reinserting the element into the blank canvas. Shuffling the final 60 bp had the same effect as shuffling the entire element, revealing that this end of the element is necessary for boundary creation (**Fig. 6d**). We then created sliding windows of 10 bp, 50 bp, and 100 bp along the element and inserted each subsequence into the blank canvas. No individual subsequence was sufficient to reproduce the effect of the entire element (**Fig. 6e**). However, shuffling the first 307 bp while maintaining the last 60 bp intact did create a strong boundary. Since the GC content of Charlie7 is unusually low, we next replaced parts of the element with random GC-matched sequence. A length-matched sequence with a GC content below 30% and the final 60 bp of the Charlie7 element was sufficient to create a boundary (**Fig. 6f**). Completely random insertions with a GC content below 30% and above 60% are also highly impactful (**Fig. S10**). Based on these *in silico* experiments, we conclude that GC content along with sequence syntax could be critical for the insulating behavior of Charlie7. Looking across all disruptive retrotransposons, we identify several families with very extreme average GC content (**Fig. S10**), suggesting the intriguing hypothesis that abrupt shifts in GC content resulting from repetitive element insertions into genomic DNA contribute to genome folding.

## Discussion

In summary, we present a whole-genome, unbiased survey of the sequence determinants of 3D genome folding using a flexible deep-learning method for scoring the effect of genetic variants on local chromatin interactions. Our study utilized synthetic mutations ranging from large deletions tiled across hundreds of megabases down to single-nucleotide perturbations within sequence motifs. Leveraging the high throughput of this *in silico* screening strategy, we showed 3D genome organization is sensitive to perturbations in A/B compartment, GC content, CTCF motif density, and active transcription. We identified clusters of transposons and RNA genes important for 3D genome folding, as modulating their sequences disrupts chromatin contacts on par with or more than modulating CTCF sites. Many of the repetitive elements with the largest effects on 3D genome folding when deleted and inserted do not contain CTCF and have not previously been implicated in chromatin architecture.

This study contributes to a growing body of evidence showing that transposable elements modulate genome folding [44] and replication timing [45]. It has long been hypothesized that transposons may have been evolutionarily co-opted as regulatory elements [35,36]. Most transposable elements are epigenetically decommissioned [46], but functional escape can change genome conformation [47]. We observe both loss and gain of contact upon transposable element deletion, supporting the idea that these elements can both establish new boundaries by installing CTCF-like motifs and inhibit ancient CTCF binding sites to block contact [37]. Our results are also consistent with previous findings that specific MIR elements and tRNAs can serve as insulators [48,49], while Alu and hAT provide loop anchors [50,51], and hint that repetitive elements may work in tandem [42]. We are curious to test coordination of transposable elements as *shadow loop anchors*, theorized by Choudhary et al. to act as redundant regulatory material supporting CTCF [37]. We anticipate that comparing disruption to element age and species divergence will help to understand the evolutionary mechanisms of transposable element deprogramming and selection in gene regulation.

Although we did not focus on CTCF specifically, a similar targeted *in silico* approach could directly address why the majority of CTCF motifs are not active [52,53], and if methylation sensitivity of CTCF motifs containing CpGs tunes folding specificity [54]. In our insertion experiments, we use a fixed and arbitrary gap between motifs. We anticipate future *in silico* experiments will refine the spacing and orientation rules of neighboring and redundant CTCF elements and reveal how CTCF coordinates with flanking proteins and transposable elements.

It is important to emphasize that our *in silico* strategy, while previously demonstrated to be highly accurate [20], is a screening and hypothesis-generating tool. Model predictions, especially those that implicate novel sequence elements or mechanisms, will require experimental validation. We view this as a strength of our approach, not a weakness. Our ability to test millions of mutations efficiently and in an unbiased manner enables us to develop hypotheses and prioritize genomic loci that would not otherwise have been considered for functional characterization. It is now a high priority to apply massively parallel reporter assays, epitope devices, and genome engineering to explore how hAT, MIR, ERV and SVA elements function in the context of 3D genome folding. We advocate for deep learning as a powerful strategy for driving experimental innovation which can be used iteratively with wet lab technologies to accelerate discovery.

Our conclusions rest heavily upon the Akita model, which only considers a limited genomic window. Future work could apply the approach presented here with other deep-learning models to test the robustness of our findings and potentially discover additional sequence features missed in our work. Our study is also limited by the quality of the hg38 reference genome, and we anticipate that extending to the new telomere-to-telomere human genome assembly will enable a better understanding of near-identical repetitive elements [33]. Finally, in order to leverage the best quality data currently available, we only made predictions for one cell type, HFFc6, but features of the 3D genome can be cell-type specific [55]. As very high-resolution and single-cell measurements of chromatin contacts, gene expression, and epigenetic states are generated for more cell types, it will be exciting to search for sequences that are necessary and sufficient for chromatin contacts in each cell type and to explore how variable these sequence determinants are across cellular contexts.

In our investigation, we develop a toolkit of *in silico* experimental strategies, including: unbiased and targeted deletion screens, phenotypic rescue, insertions into synthetic sequence, sampling around known sequence motifs, and sequence contribution tracks across tens of basepairs to megabases. We hope that the variety of experiments profiled here may serve as a template for foundational biological research with deep learning. We also anticipate that our released disruption tracks will provide useful annotations for genome sensitivity and yield further insights into chromatin biology. In sum, our work highlights the potential of deep learning models as powerful tools for biological hypothesis generation and discovery in regulatory genomics.

## Methods

### Akita model and datasets

Throughout this analysis, we use the published convolutional neural network Akita to predict log(observed/expected) chromatin contact maps from ∼1 Mb (1,048,576 bp) of real, altered, or synthetic DNA sequence[20] (https://github.com/calico/basenji/tree/master/manuscripts/akita). All types of mutations, including deletions, insertions, inversions and substitutions, may be scored as long as they are smaller than 1 Mb. Akita’s predictions have been shown to mirror experimental results with deletions across scales of thousands of base pairs (bp) to single nucleotides. Fudenberg et al. originally trained Akita across six cell-types simultaneously, and we made all predictions in this work in the cell-type with the highest resolution of training data, human foreskin fibroblasts (HFFc6). The experimental Micro-C maps from HFFc6 [28] are used in visualizations. All epigenetic and transcriptomic data were generated in HFFc6 and downloaded from public repositories. The source of all public data, including Micro-C, ATAC-Seq, RNA-Seq, ChIP-Seq, and compartment calls, can be found in Data Table 1. All analyses use the hg38 genome build. We downloaded centromere locations from UCSC Table Browser [56].

## Data Table 1

| Datatype | Source | Accession No. |
| --- | --- | --- |
| Micro-C, HFF | 4DN, Krietenstein et al. 2020[28] | 4DNESWST3UBH |
| RNA-Seq, HFF | ENCODE [29], GEO:GSE188028 | ENCFF262XNE, ENCFF381OAF |
| ChIP-Seq, CTCF, HFF | ENCODE, Lou et al., 2020[57] | ENCFF465OQP |
| ATAC-Seq, HFF | 4DN, Oksuz et al., 2021[58] | 4DNESMBA9T3L |
| Compartment, HFF | 4DN, Oksuz et al., 2021[58] | 4DNES9X112GZ |

### Pipeline for computing 3D genome folding disruption scores

The location of deletions and insertions are centered such that the start position of the variant is always introduced halfway through the 1-Mb sequence at 2^19^ bp. For deletions, we pull additional sequence from the right to pad the input to 2^20^ bp. We remove sequences from our analysis which overlap centromeres [59], ENCODE blacklisted regions [60], and regions with an N content greater than 5%. Evaluating predictions on GPU (NVIDIA GeForce GTX 1080 Ti, NVIDIA TITAN Xp, NVIDIA GeForce RTX 2080 Ti) decreased the time per variant from 1.58 seconds to 262 ms, on average.

We score disruption as the log of the mean squared error between reference and perturbed maps. Mean squared error captures large-scale contact map changes, and has been used previously to rank predictions [20]. Pearson/Spearman correlation is also an appropriate choice [27].

### Mass deletion screens

Along with controls, we perform the following large-scale deletion screens:

1. *5 kb, whole genome* (*n* = 562,743).
2. *10,000 (10k) random CTCF deletions*. CTCF locations are pulled from JASPAR 2022 [61].
3. *10k 100-bp random deletions*. Start locations are randomly sampled from the genome.
4. *Randomly sized deletions*, ranging from 1 bp to 100 kb (*n* = 41,207). Start locations are randomly sampled from the genome.
5. *RepeatMasker database deletions* (*n* = 1,164,107) *[62]*. RepeatMasker downloaded from UCSC Table Browser. We exclude ambiguous elements (containing ‘?’ in the label). We initially sample 10,000 elements per family or up to the total number of elements in the family, whichever is less. Thereafter, we randomly sample from the database.
6. *TSS deletions*. (*n* = 1,073,329 mutations across 1,789 genes).

A full summary as well as the location of these results can be found in **Supplementary Table 1**.

### Genomic tracks

We smoothed the disruption scores of 5-kb deletions with a rolling average of 50 bp to create disruption tracks (**Fig. 1d, Fig. 3a**). We additionally visualize the density of the following elements at 5-kb resolution:

1. Reference genes, hg38, GENCODE v39 [63], downloaded from UCSC Table Browser.
2. ENCODE hg38 v3 candidate cCREs, ENCODE Project [29], downloaded from UCSC Table Browser.
3. CTCF motifs (MA0139.1), JASPAR 2022 [61], downloaded from http://expdata.cmmt.ubc.ca/JASPAR/downloads/UCSC_tracks/2022/hg38/.
4. ATAC-Seq peaks in HFFc6 [58].
5. Alu, L1, and L2 elements, RepeatMasker database, v. 4.1.2 [62], downloaded from UCSC Table Browser.

### Overlap with compartment, ENCODE cCREs, CTCF, and transcribed genes

We used pre-computed compartment scores generated from the HFFc6 Micro-C dataset originally employed for training Akita [28]. To calculate the overlap between disruption scores for 5-kb deletions and compartment scores generated at 50-kb resolution, we merged both measures by genomic location, filled missing disruption values with linear interpolation, and calculated the overlap across A compartments with a compartment score greater than 0 and B compartments with a compartment score less than 0.

We intersected deleted windows and transposable elements with ENCODE cCREs using bioframe [64] to calculate the percentage overlap. We use the same strategy to calculate overlap with JASPAR CTCF motifs, ATAC-Seq peaks, and transcribed elements. When quantifying transcription of repetitive elements unannotated as genes, we calculated overlap with RNA-seq BigWigs, summed across both strands.

### Mappability

Per nucleotide mappability was measured using 24-kmer multi-read mappability, where mappability is the probability that a randomly selected read of length k in a given region is uniquely mappable [65]. Mappability tracks were downloaded from the Hoffman lab (https://bismap.hoffmanlab.org). In this study, mappability averaged across 5 kb deletions, repetitive element families, and Alu element types in a 100 Mb subset of chromosome 1 from 100 Mb to 200 Mb.

### *In silico* mutagenesis at the TSS

We examined behavior at the TSS using *in silico* mutagenesis. We individually randomly mutated each nucleotide 300 bp upstream to 300 bp downstream of the top 1,789 highest expressed protein coding genes via total RNA-Seq and quantified the MSE between mutated and reference predicted maps. We observed that 1,015 genes fell in A compartments, while 63 fell in B compartments. To produce tracks in **Fig. 2f-g**, we averaged the disruption of each nucleotide by position and smoothed using a rolling average of 20 bp. We used the same strategy across select repetitive elements to identify which nucleotides most contribute to entire-element disruption scores (**Fig. 5e**). To create metaplots, we selected the highest scoring nucleotide change for each gene, and filtered all genes with a maximum disruption score above -7. We then averaged the difference between reference and perturbed maps for these genes.

### Repetitive elements

Repetitive element density was calculated as the number of elements across the entire RepeatMasker database overlapping each 5-kb genomic bin. We quantified enrichment as the log fold change of the mean disruption across 10% of genomic windows per family compared to all windows. To create metaplots, we average the difference between maps for the top 100 repetitive element deletions per family, along with CTCF deletions.

### Phenotypic Rescue

We profiled the following elements in our proof-of-concept phenotype rescue screen:

1. A MER91B hAT-Tip100 element at position chr2:98412915-98413053. SWA score: 392, Divergence: 27%. Disruption from reference = -2.65.
2. A size-matched 138-bp random DNA sequence. Disruption from deletion = -2.55.
3. The canonical CTCF motif (TGGCCACCAGGGGGCGCTA). Disruption = -2.68.
4. A MER91B element at position chr12:51824097-51824219.

SWA score: 245, Divergence: 20.9%. Disruption = -5.28.

### Insertion Screens

#### CTCF depletion

We created a simulated Hi-C contact map without structure as a blank canvas for insertion experiments. We first generated a random DNA sequence of length 2^20^ bp. By chance, predicted maps from random sequence will contain some above background contact frequencies. To remove all structure, we incremented across this sequence one nucleotide at a time with a 12-bp sliding window. For each position, we computed the edit distance to the consensus CTCF motif. If the edit distance fell below a set threshold, we inserted a random DNA sequence of length 12 until the subsequence was sufficiently different from CTCF. Experimenting with edit distances, we found that a distance of 7 produces predicted maps which lack structure but do not result in artificial model predictions (**Fig. S6**). We call this a “blank canvas” 1-Mb sequence.

#### CTCF insertion

We inserted the CTCF motif into the blank canvas and predicted expected contact frequencies with Akita. We quantify insertion impact as the log mean squared error between the predicted maps of the blank canvas and the insertion. If more than one motif was added, the insertions were centered and separated by an arbitrary 100 bp. To sample the CTCF motif, we drew frequencies from the CTCF position weight matrix [61]. To create a baseline, we inserted 5,000 CTCF motifs drawn from locations in the genome. Sequence motifs were visualized with a python port of the seqLogo package [66,67].

#### Repetitive element insertions

We selected the top 1,000 most disruptive repetitive elements per family by the deletion screen to insert back into the blank canvas sequence. We inserted both the forward and reverse complement of each sequence, and selected the direction with the highest score. For an initial screen, we inserted all elements 100x with 100-bp spacing. As an additional baseline, we inserted 1,000 201-bp randomly generated sequences, as the median repetitive element size in our insertion screen was 201 bp. To perform clustering with t-SNE, we decreased the resolution of the 448×448 pixel maps to 100×100 pixels and flatten them to 1D vectors before clustering.

#### Additional genomic tracks

In **Fig. 5e**, we visualized CTCF ChIP-Seq and CTCF motif locations in the element’s original genomic context. Along with deleting the entire element, we performed mutagenesis to a random nucleotide across the length of the element to create a ‘disruption track’ of nucleotides most sensitive to perturbation. We highlight the most sensitive bases.

#### JASPAR Insertions

We inserted the forward and reverse complement of each JASPAR motif [61] into CTCF-depleted sequence with 100-bp spacing (*n* = 842). JASPAR motifs were pulled and coordinated with pyJASPAR [68].

### Code and Data Availability

Disruption results are available at: github.com/keiserlab/3d-genome-disruption-paper

### Supplemental Note

One concern is sequence mappability potentially confounding model training. Repetitive elements are, by nature, highly conserved and present inherent difficulties assigning multi-mapped reads. Before training the model, large gaps were excluded from the training dataset and missing Hi-C bins were linearly interpolated [20]. If repetitive elements were systematically removed or imputed, the model may behave unreliably when predicting unseen repetitive element sequences.

To investigate this confounder, we examined how sequence mappability compares to disruption score (**Fig. S4)**. In general, we observe no correlation between deletions of 5-kb windows and mappability, indicating that poorly mappable sequences do not have unusually high or low disruption scores. Mappability of individual elements is also uncorrelated with disruption.

We do find that Alu elements have particularly low sequence mappability and particularly high predicted importance. Many Alu elements are still active and recently inserted into DNA, and therefore have high sequence similarly, presenting a challenge in mapping. It is also possible that the highly conserved nature of recent Alu elements contributes to their utility in shaping the 3D genome. The correlation with mappability is expected and may or may not indicate a bias; it is difficult to disentangle these two possibilities easily. Relatively low negative correlation between disruption score and mappability for individual elements within the Alu class suggests that many of the highly disruptive Alus are not in regions of low mappability.

## Acknowledgements

We gratefully acknowledge Vijay Ramani, Elphege Nora, Tony Capra, and Geoff Fudenberg for valuable scientific insights and project guidance. We thank Evonne McArthur for generous discussion and the idea to insert motifs into CTCF-depleted random sequence. We additionally thank members of the Pollard and Keiser labs for useful comments and manuscript feedback.

## Author Contributions

L.M.G., K.S.P., and M.J.K. conceptualized the work, designed experiments and guided the project direction. L.M.G. conducted all analyses and prepared figures, with feedback from K.S.P. and M.J.K. L.M.G. and K.S.P. drafted the original manuscript. L.M.G., K.S.P., and M.J.K. reviewed and edited the manuscript. K.S.P. and M.J.K. acquired funding.

## Funding Sources

This work was supported by the NIH 4D Nucleome Project (grant #U01HL157989 to K.S.P.), grant number 2018-191905 from the Chan Zuckerberg Initiative DAF (M.J.K.), an advised fund of the Silicon Valley Community Foundation (M.J.K.), and a UCSF Achievement Rewards for College Scientists (ARCS) Scholarship (L.M.G.).

## Supplementary Figures

**Supplementary Figure 1:**
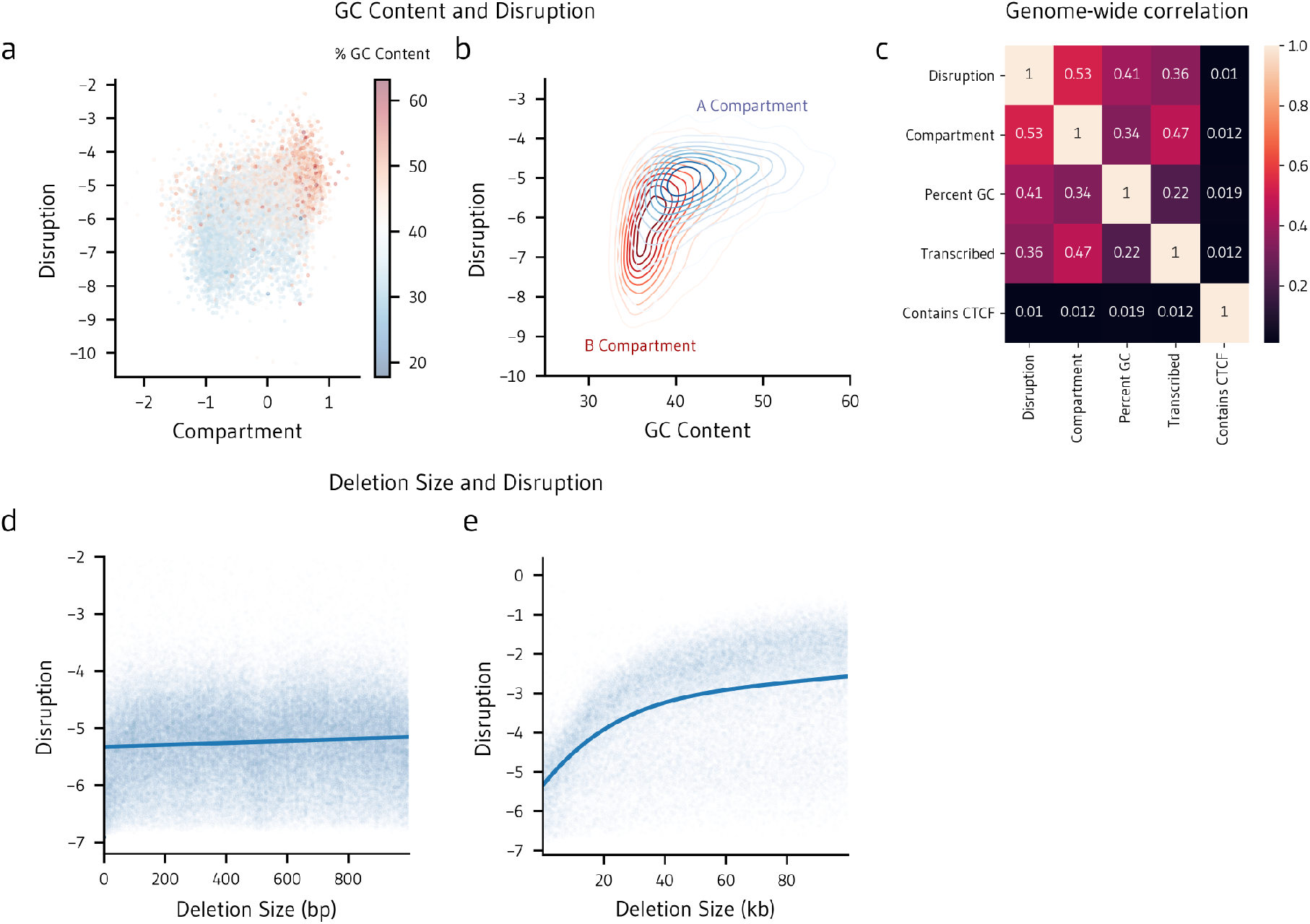
Disruption is correlated with GC content and deletion size. **a**. Disruption scores across the 5 kb whole-genome deletion screen compared to compartment score, as defined as the first eigenvector of the experimental micro-C contact matrix in HFFc6. **b**. GC Content across the 5-kb screen compared to disruption score. **c**. Disruption scores across a deletion screen of random sized genomic segments ranging from 1 bp to 1,000 bp across chromosome 17 (*n* = 2,000). **d**. Disruption scores across a deletion screen of random sizes ranging from 1 bp to 100,000 bp across chromosome 1 (*n* = 39,207).

**Supplementary Figure 2:**
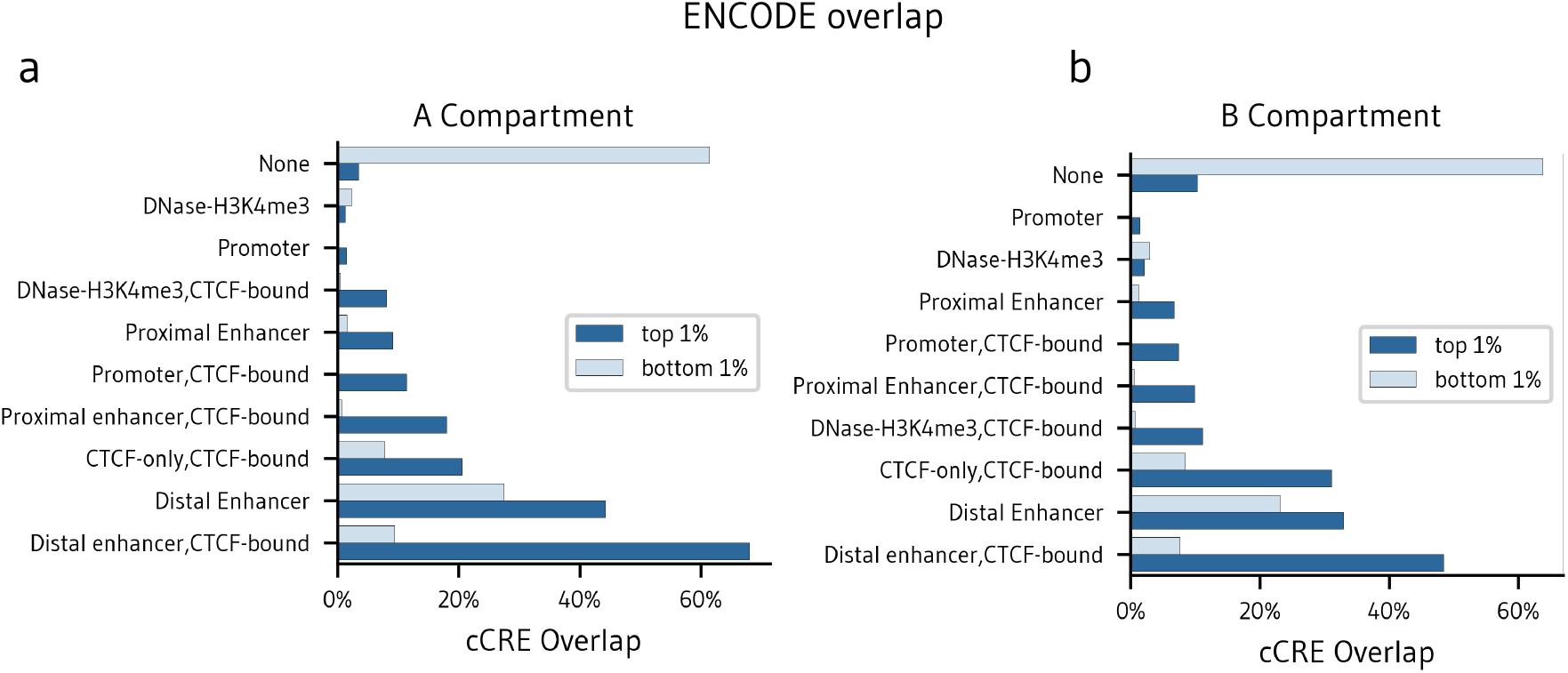
CTCF Enrichment in A and B Compartments. CTCF-bound regions are enriched within the top 1% most disruptive 5 kb regions compared to the bottom 1% in both A compartments (**a**) and B compartments (**b**).

**Supplementary Figure 3:**
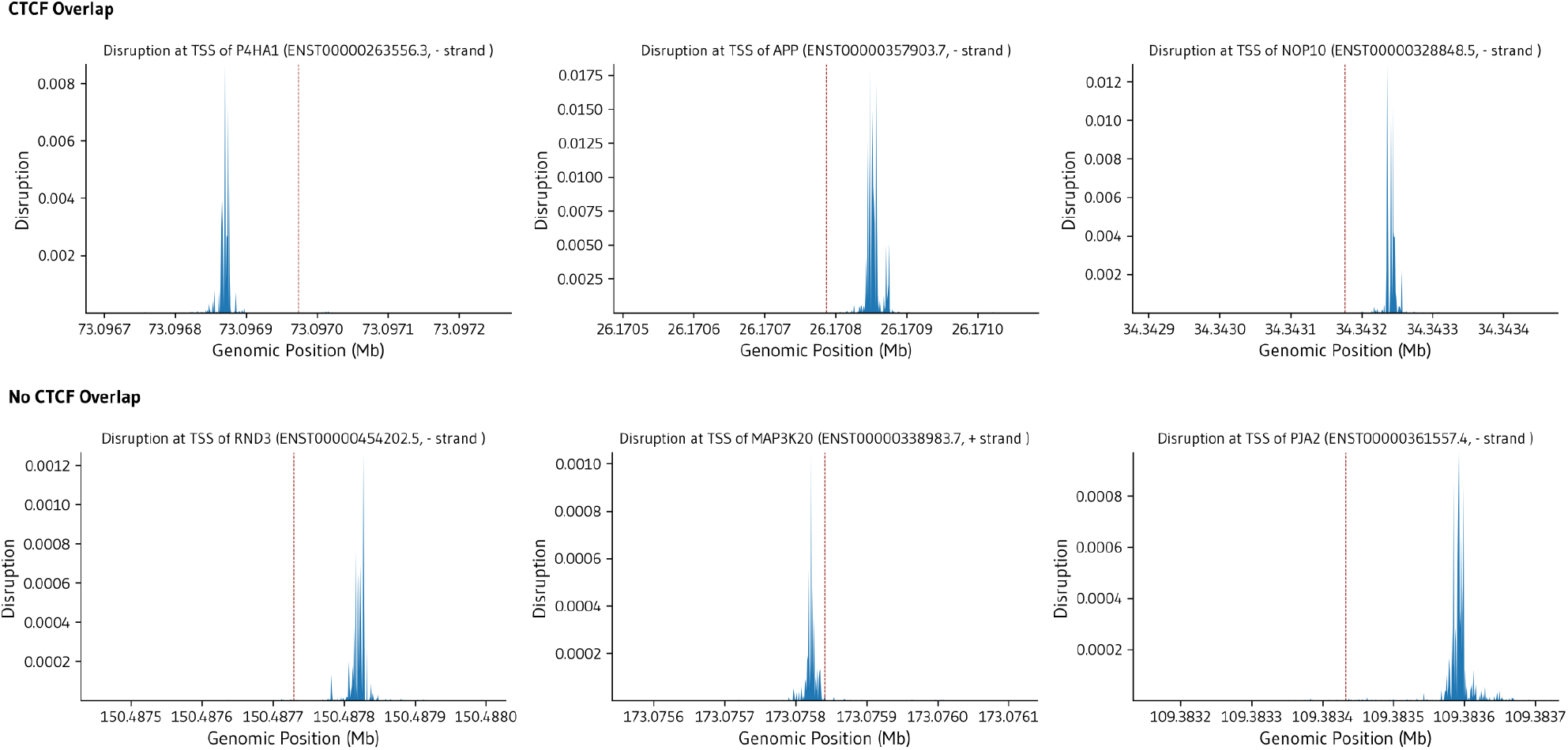
Transcription tracks. Individual single-nucleotide disruption tracks around the TSS of highly expressed genes which overlap CTCF (top) and do not overlap CTCF (bottom). The location of the TSS is marked in red.

**Supplementary Figure 4:**
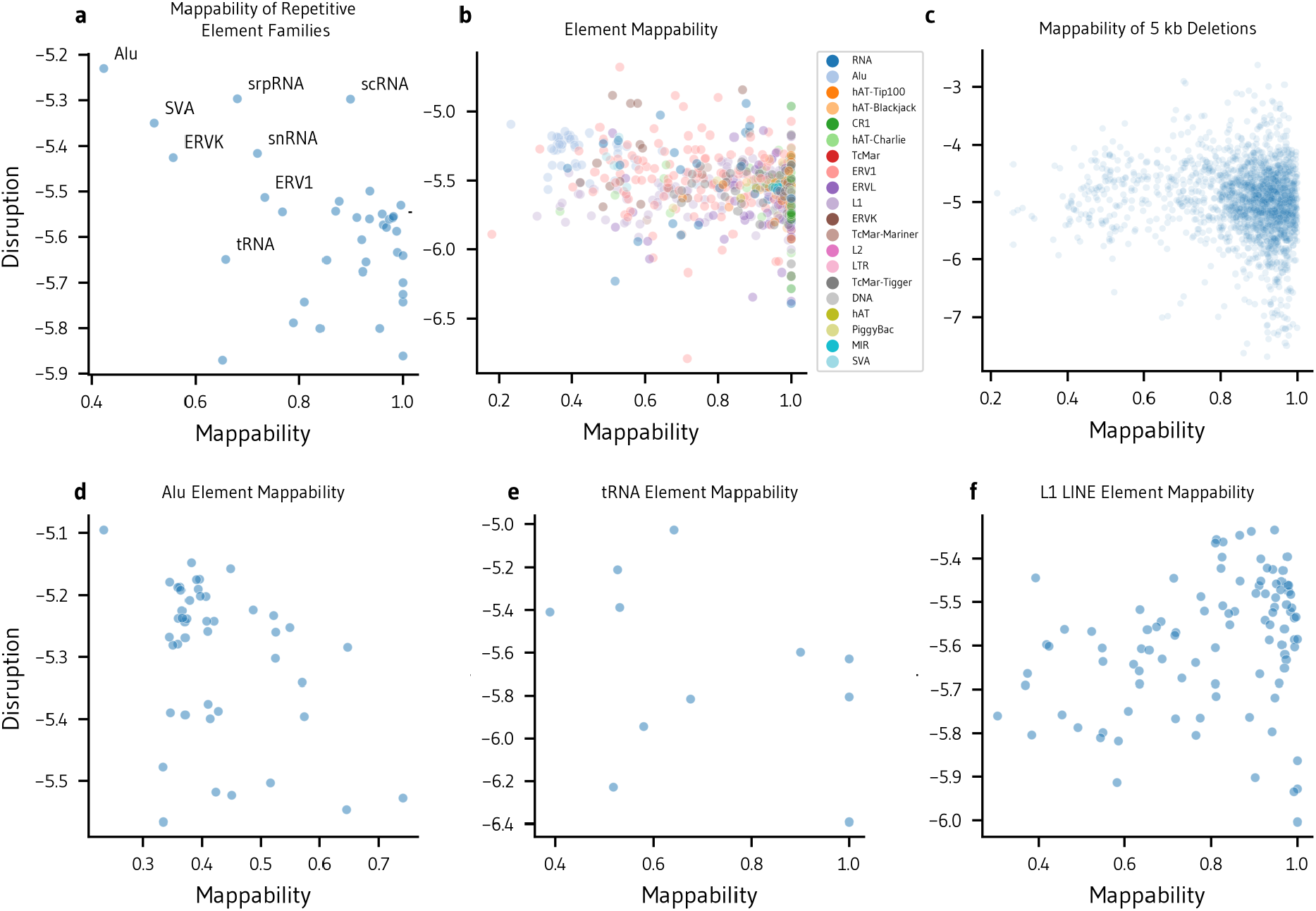
Disruption and Mappability. Comparison of multi-read mappability at chr1:100Mb-200Mb and disruption scores. **a**. Average mappability by repetitive element family. **b**. Average mappability by repetitive element type. **c**. Average mappability of 5-kb deleted genome windows. **d-f**. Average mappability of repetitive element types within the Alu, tRNA, and L1 LINE families.

**Supplementary Figure 5:**
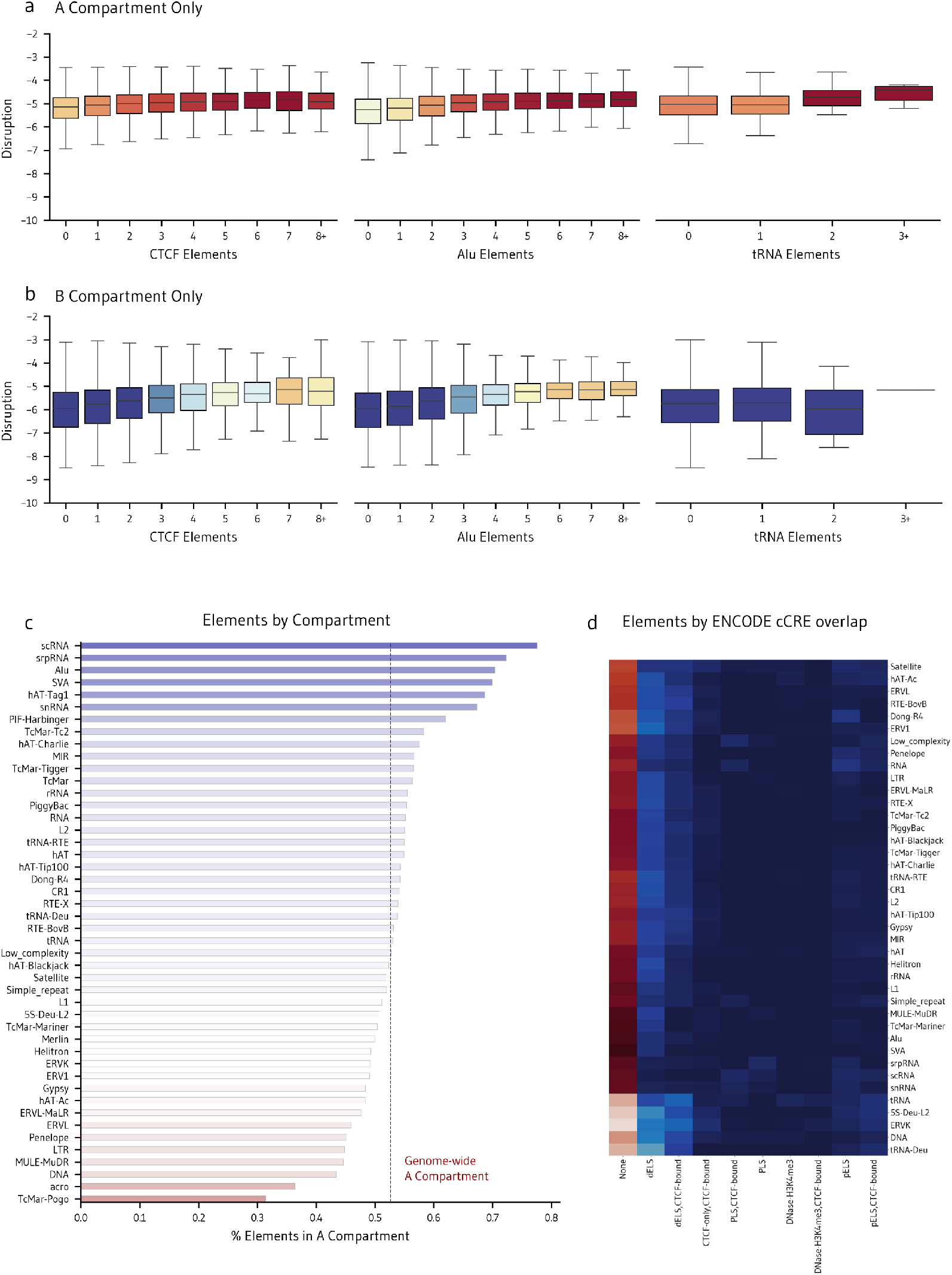
Repetitive elements vary across compartment and regulatory region. **a-b**. Disruption by repetitive element count within 5 kb genomic windows, by compartment. **c**. Percent of each sampled repetitive element family found within the A compartment, as defined by the first eigenvector of experimental micro-C in HFFc6. **d**. Overlap of top 10% most disruptive repetitive elements by the deletion screen stratified by overlap with ENCODE candidate regulatory elements (cCREs).

**Supplementary Figure 6:**
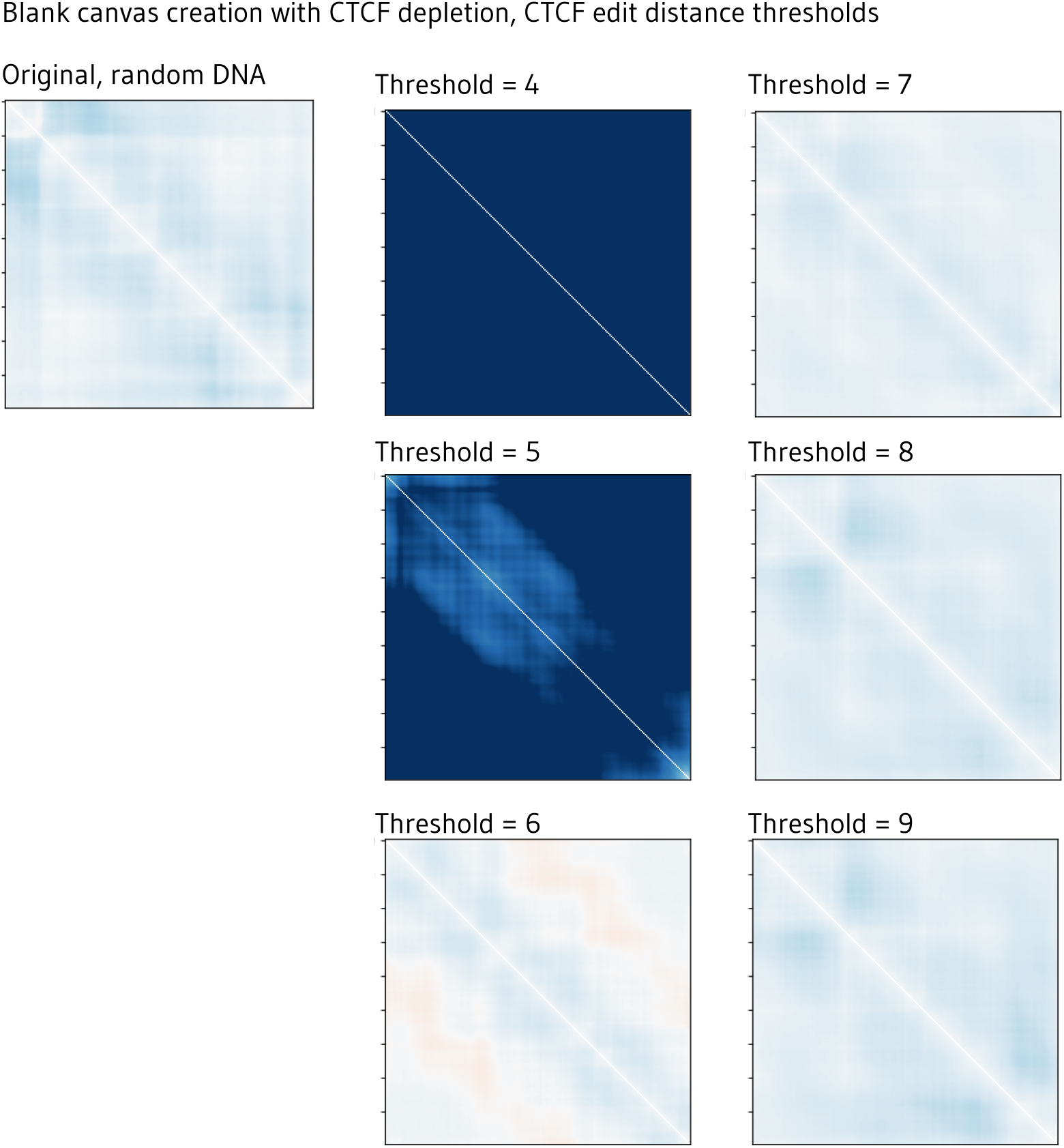
Edit distance thresholds for blank canvas map creation. We create a blank map to insert elements by predicting genome folding of random DNA sequence. The original map, by chance, contains spurious structure, so we deplete the sequence of any subsequence within a given edit distance of CTCF. An aggressive threshold (e.g. 4) does not produce a biologically plausible sequence, while a permissive threshold (e.g. 9) leaves structure. We select a threshold of 7.

**Supplementary Figure 7:**
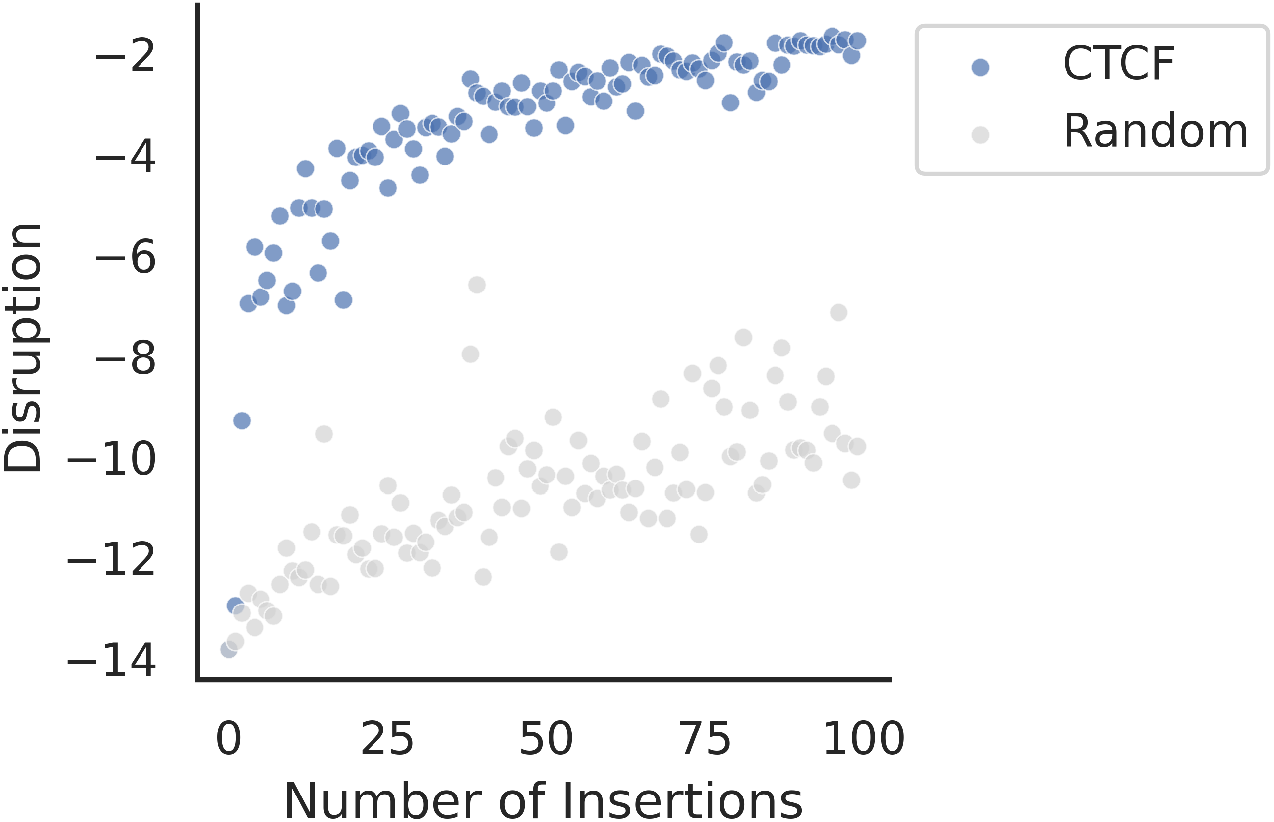
Motif insertion strength. Impact of increasing the number of CTCF and random 12-bp sequence insertions into a blank map. Insertions are separated by 100 bp randomly generated DNA sequence.

**Supplementary Figure 8:**
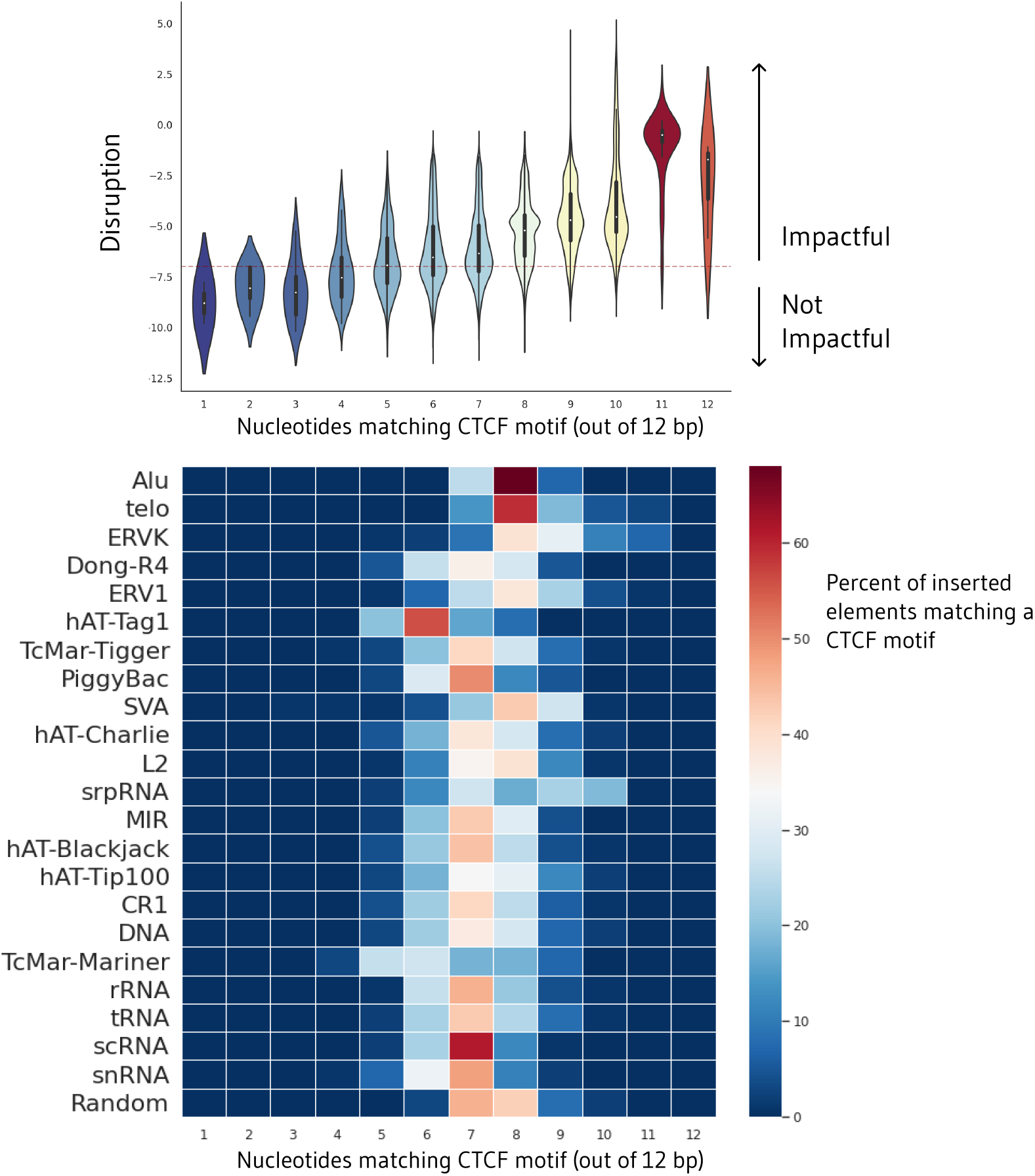
Similarity of impactful insertions to CTCF. Number of nucleotides of inserted repetitive elements matching the consensus CTCF motif versus element disruption score (top). Elements were scanned one nucleotide at a time to calculate the edit distance of all 12-bp subsequences to CTCF. 12 indicates that the element contains a perfect CTCF motif match. 1 indicates the element contains no subsequences matching the CTCF motif. Motifs more similar to CTCF are higher scoring. Number of nucleotides of inserted repetitive elements matching the CTCF motif, by family (bottom). Only elements with a disruption score above -7 (red threshold) are shown below.

**Supplementary Figure 9:**
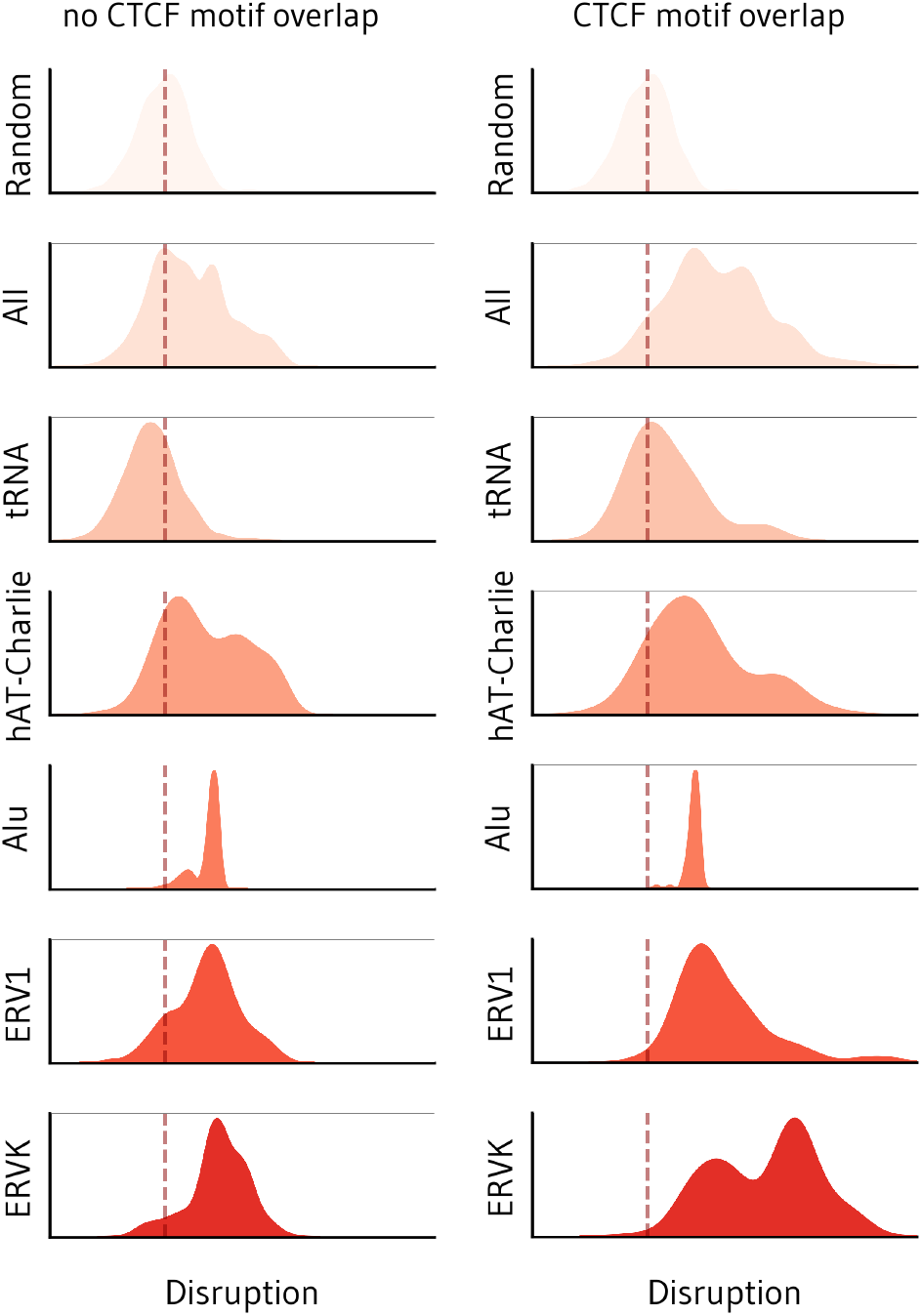
Impact of repetitive element insertions with and without CTCF. Disruption scores across all repetitive element insertions into a blank, CTCF-depleted map, striated by overlap of the original element with an annotated JASPAR CTCF motif.

**Supplementary Figure 10:**
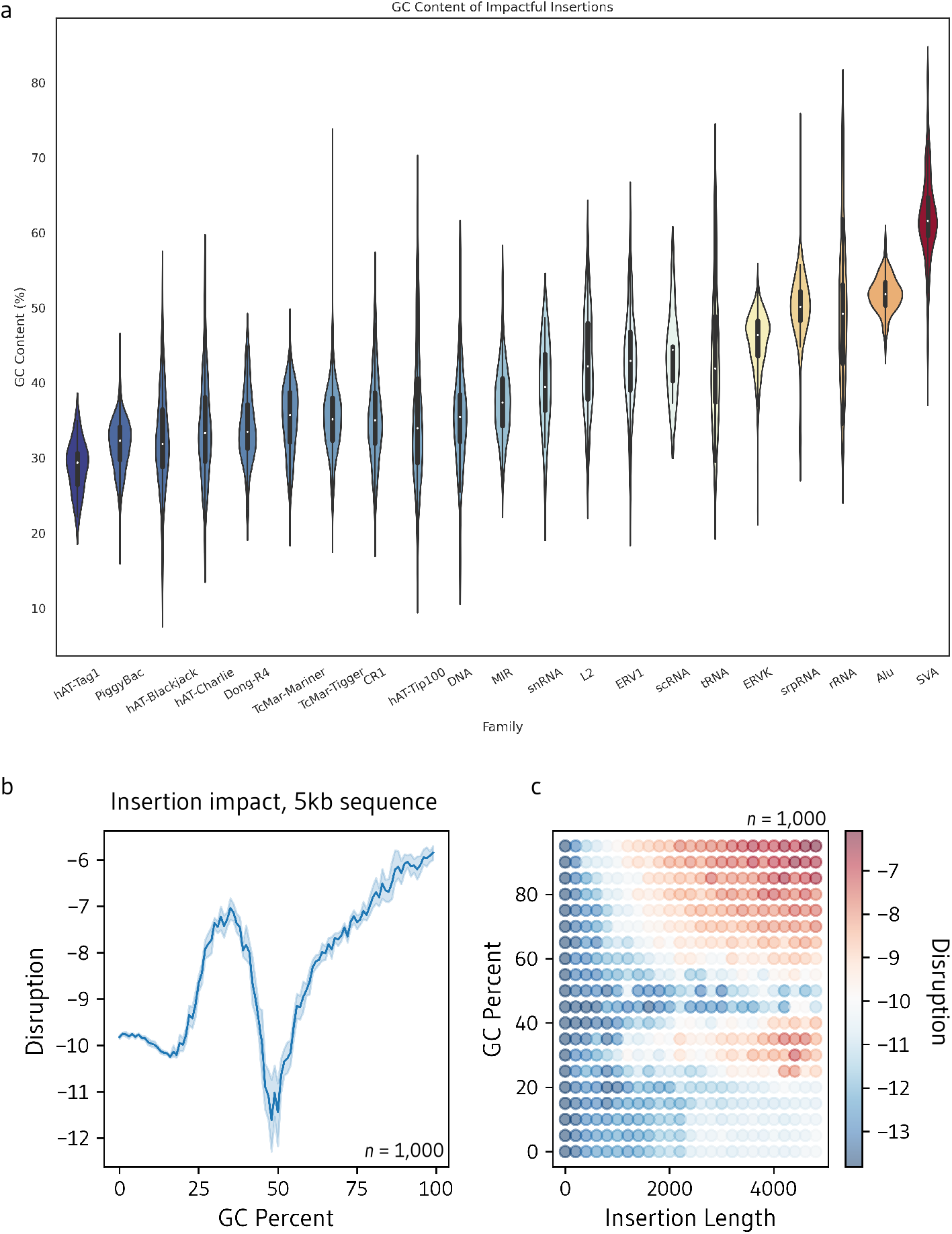
GC content of impactful insertions. **a**. GC content of disruptive repetitive elements with a MSE greater than -5 upon insertion into a blank map. **b**. Disruption caused by insertion of randomly generated 5-kb DNA sequences with GC percentages ranging from 0% to 100%. **c**. Disruption produced by random insertions into a blank map ranging from GC percentages from 0% to 100% and lengths from 1 bp to 5 kb.

**Supplementary Data Table 1.**

| Filename | n | Type | Context | Description |
| --- | --- | --- | --- | --- |
| 5kb_deletions.csv | 562,744 | deletion | original | Tiled 5kb deletions across the genome. |
| random_sized_deletions_100kb.csv | 39,207 | deletions | original | Random deletions between 1 and 100kb from chromosome 1. |
| random_sized_deletions_1kb.csv | 2,000 | deletions | original | Random deletions between 1 and 1kb from chromosome 17. |
| RepeatMasker_deletions.csv | 1,164,108 | deletions | original | Deletions from the RepeatMasker database. |
| CTCF_motif_insertions.csv | 991 | insertion | blank | Sampled CTCF JASPAR motifs inserted into blank map. |
| CTCF_PWM_sampling_insertions.csv | 326,177 | insertion | blank | Sampled CTCF PWM inserted into blank map. |
| random_20bp_insertions.csv | 236,119 | insertion | blank | Random 20bp sequences inserted into blank map. |
| CTCF_consensus_motif_insertions.csv | 1000 | insertion | blank | CTCF consensus sequence inserted into blank map (different random sequence spacing). |
| random_201bp_insertions.csv | 1000 | insertion | blank | Random DNA sequence inserted into blank map. |
| all_motif_insertions.csv | 842 | insertion | blank | JASPAR motif insertions into blank map. |
| RepeatMasker_insertions.csv | 14514 | insertion | blank | RepeatMasker element insertions into blank map. |
| tss_mutagenesis.csv | 1,073,334 | mutagenesis | original | Random single nucleotide mutagenesis around the TSS. |

## Notes

### Competing Interest Statement

The authors have declared no competing interest.

